# Quantifying Differences and Similarities in Whole-brain White Matter Architecture Using Local Connectome Fingerprints

**DOI:** 10.1101/043778

**Authors:** Fang-Cheng Yeh, Jean M. Vettel, Aarti Singh, Barnabas Poczos, Scott Grafton, Kirk I. Erickson, Wen-Yih I. Tseng, Timothy D. Verstynen

## Abstract

Quantifying differences or similarities in connectomes has been a challenge due to the immense complexity of global brain networks. Here we introduce a noninvasive method that uses diffusion MRI to characterize whole-brain white matter architecture as a single local connectome fingerprint that allows for a direct comparison between structural connectomes. In four independently acquired data sets with repeated scans (total N=213), we show that the local connectome fingerprint is highly specific to an individual, allowing for an accurate self-versus-others classification that achieved 100% accuracy across 17,398 identification tests. The estimated classification error was approximately one thousand times smaller than fingerprints derived from diffusivity-based measures or region-to-region connectivity patterns. The local connectome fingerprint also revealed neuroplasticity within an individual reflected as a decreasing trend in self-similarity across time, whereas this change was not observed in the diffusivity measures. Moreover, the local connectome fingerprint can be used as a phenotypic marker, revealing 12.51% similarity between monozygotic twins, 5.14% between dizygotic twins, and 4.51% between none-twin siblings. This novel approach opens a new door for probing the influence of pathological, genetic, social, or environmental factors on the unique configuration of the human connectome.

**Author Summary:** The local organization of white matter architecture is highly unique to individuals, making it a tangible metric of connectomic differences. The variability in local white matter architecture is found to be partially determined by genetic factors, but largely plastic across time. This approach opens a new door for probing the influence of pathological, genetic, social, or environmental factors on the unique configuration of the human connectome.

## Introduction

The specific brain characteristics that define an individual are encoded by the unique pattern of connections between the billions of neurons in the brain [1]. This complex wiring system, termed the connectome [2, 3], reflects the specific architecture of region-to-region connectivity [4] that supports nearly all complex brain functions. Yet to date, quantifying the difference between connectomes of two or more individuals remains a major challenge, as it requires a reliable characterization of white matter architecture that is also sensitive to microscopic variability.

To this end, studies have used diffusion MRI (dMRI) to measure the architecture of white matter pathways using the diffusion properties of water molecules[5, 6]. This allows for the mapping of white matter trajectories in the human brain and defining the graph structure of region-to-region connectivity [7, 8]; however, while the reliability of diffusion MRI scans has improved substantially by new acquisition approaches [9, 10], the efficiency and accuracy of tractography approaches have recently come into question [11, 12]. Thus, instead of mapping region-to-region connectivity, the concept of the *local* connectome was proposed as an alternative measure to overcome the limitations of diffusion MRI fiber tracking [11-13]. The local connectome is defined as the degree of connectivity between adjacent voxels within a white matter fascicle measured by the density of the diffusing water. A collection of these density measurements provides a high dimensional feature vector that can describe the unique configuration of the structural connectome within an individual, providing a novel approach for comparing differences and similarities between individuals as pairwise distances.

In this study, we used this local connectome feature vector as a fingerprint to quantify similarities and difference between two white matter architectures. To evaluate the performance of our approach, we used four independently collected dMRI datasets (*n*=11, 25, 60, 118, see Methods) with repeat scans at different time intervals (ranging from the same day to a year) to examine whether local connectome fingerprints can reliably distinguish the difference between within-subject and between-subject scans. We then examined whether the local connectome fingerprint is a unique identifier of an individual person by testing whether the fingerprint could determine if two samples came from the same person or different individuals. This uniqueness was compared with fingerprints derived from fractional anisotropy (FA)[14], diffusivities, and conventional region-to-region connectivity methods. Follow-up analysis revealed how local connectome fingerprints can quantify the similarity between genetically related individuals as well as measure longitudinal changes within an individual.

## Results

### Characterization of white matter architecture

We first illustrate how the local connectome fingerprint uses the density of diffusing spins to characterize white matter architecture within an individual. Figure 1A shows the spin distribution functions (SDFs) [15] estimated from dMRI scans at the mid-sagittal section of the corpus callosum. SDF measures the density of water diffusing at any orientation within a voxel and the SDF magnitude at the fiber directions can quantify the connectivity of local connectome (see Methods). An example of the local connectome quantified at the corpus callosum is illustrated for three subjects in Fig. 1B. Here the anterior and posterior portion of corpus callosum exhibit substantial diversity between these three subjects. A repeat scan several months later reveals a qualitative within-subject consistency. This high individuality appears to be specific to diffusion density estimates. Conventional FA measures calculated from diffusivity do not yield this qualitative between-subject diversity (Fig. 1C).

To sample the local density measurements across all major white matter pathways, dMRI data was reconstructed into a standard space, and the fiber directions of a common atlas were used to sample an SDF value for each fiber direction (see Methods and Fig. 2A). This approach yielded, for each dMRI scan, a local connectome fingerprint consisting of a high-dimensional feature vector with a total of 513,316 density estimates (Fig. 2B). Fig. 2C shows the fingerprints of the same three subjects in Fig. 1B and the fingerprints from their repeat scans. Consistent with the qualitative measurements in Fig. 1B, each local connectome fingerprint in Fig. 2C shows, at a coarse level, a highly similar pattern for within-subject scans and also high variability across subjects, suggesting that the local connectome fingerprint may exhibit the unique features of the white matter architecture.

**Fig 1.**
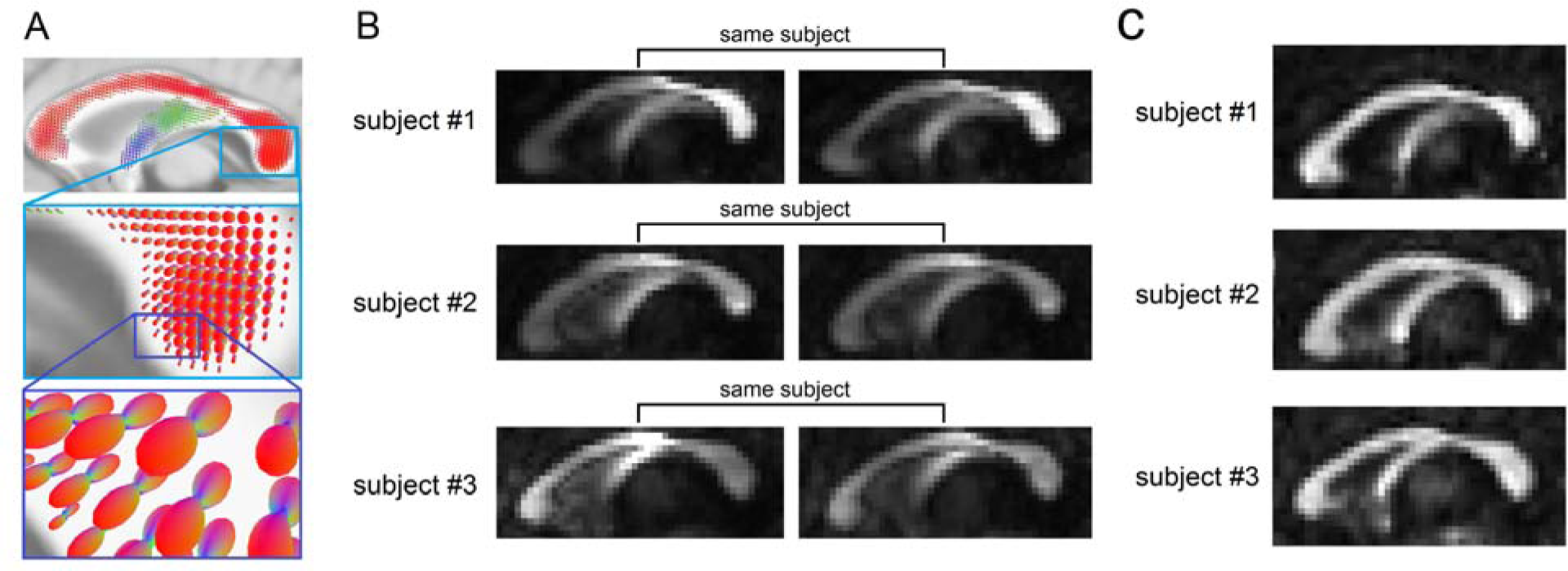
The uniqueness of local connectome structure revealed by the density of diffusing water. (A) The spin distribution function (SDF) calculated from diffusion MRI quantifies the density of diffusing water along axonal fiber bundles. The magnitudes of the SDF at axonal directions provide density-based measurements to characterize axonal fiber bundles. (B) The density measurements obtained from the SDFs show individuality between-subjects #1, #2, and #3 (intensity scaled between [0 0.8]). The density of diffusing water varies substantially across different portions of the corpus callosum. The repeat measurements after 238 (subject #1), 191 (subject #2), and 198 (subject #3) days present a consistent pattern that captures individual variability. (C) In contrast to the SDF shown in (B), the fractional anisotropy derived from diffusivity shows no obvious individuality between the same subjects #1, #2, and #3 (intensity also scaled between [0 0.8]). This is due to the fact that diffusivity, which quantifies how *fast* water diffuses, does not vary a lot in normal axonal bundles.

**Fig 2.**
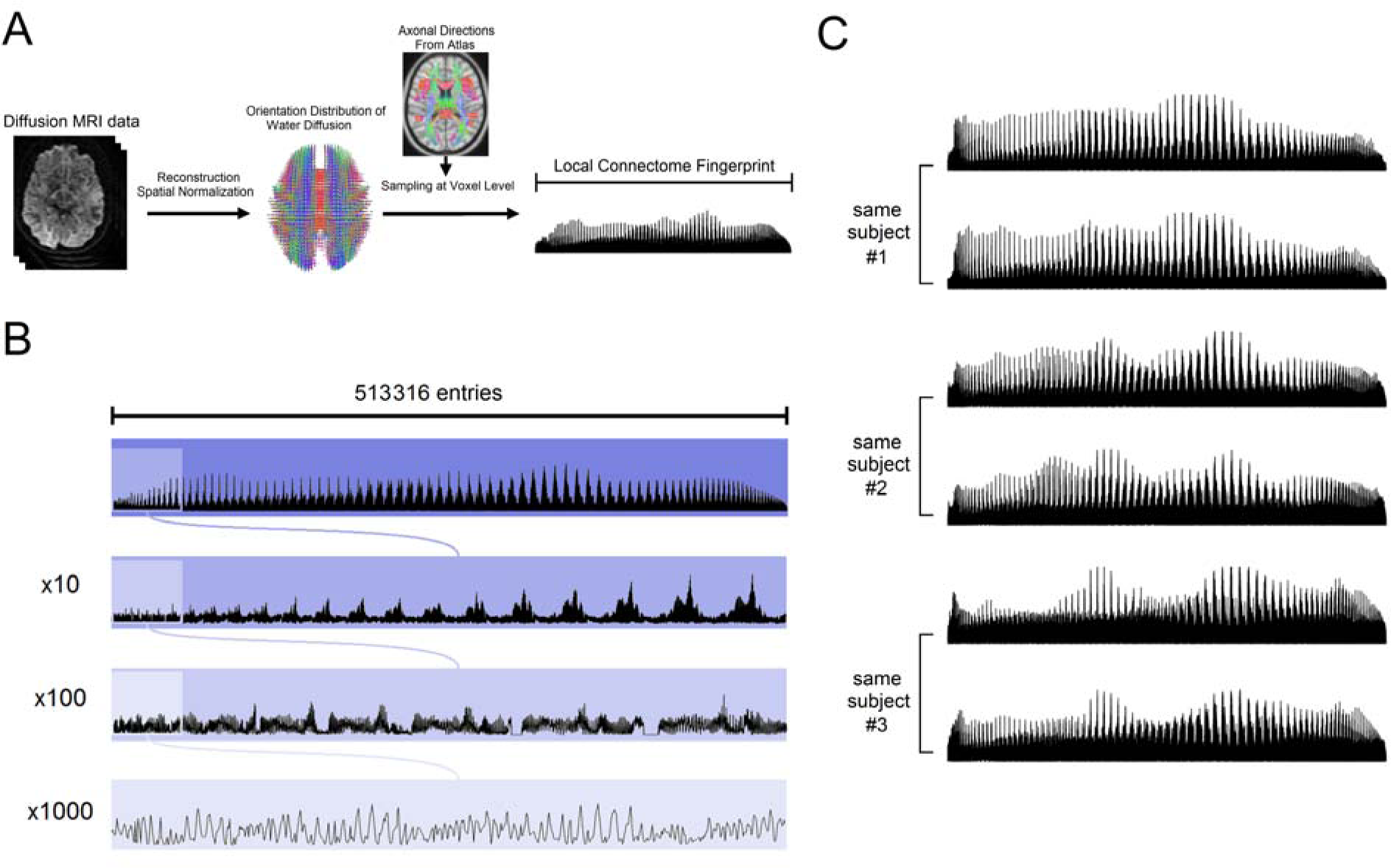
Local connectome fingerprinting. (A) Local connectome fingerprinting is conducted by first reconstructing diffusion MRI data into a standard space to calculate the spin distribution functions (SDFs). A common fiber direction atlas is then used to sample the density of diffusing water along the fiber directions in the cerebral white matter. The sampled measurements are compiled in a left-posterior-superior order to form a sequence of characteristic values as the local connectome fingerprint. (B) One local connectome fingerprint is shown in different zoom-in resolutions. A local connectome fingerprint has a total of 513,316 entries of scalar values. (C) The local connectome fingerprint of subject #1, #2, and #3 and their repeat measurements (lower row) after 238, 191, and 198 days, respectively. At a coarse level, the local connectome fingerprint differs substantially between three subjects, whereas those from the repeat scans show a remarkably identical pattern, indicating the uniqueness and reproducibility of the local connectome fingerprint.

### Between-subject versus within-subject difference

To quantify how well the local connectome fingerprint captures between-subject difference, we used four independently collected dMRI datasets (*n*=11, 24, 60, 118) with repeat scans for a subset of the subjects (*n*=11×3, 24×2, 14×2, 44×2, respectively). The Euclidian difference (i.e., root-mean-squared error) was used as a single difference estimate between any two fingerprints. For each dataset, we computed within-subject differences (n=33, 24, 14, 44, respectively) and between-subject differences (n=495, 1104, 2687, 12997, respectively). Figure 3 shows the within-subject and between-subject differences of the four datasets. All four datasets show a clear separation between the within-subject and between-subject difference distributions, with no single within-subject pair as large as any of the between-subject pairs. We used d-prime [16] to quantify the separation of between-subject and within-subject differences. The results showed d-prime values of 14.84, 12.80, 7.21, and 8.12, for dataset I, II, III, and IV respectively, suggesting a very high degree of separation between the two distributions.

In order to understand what regions of the local connectome may be driving this within-subject uniqueness, we looked at the spatial distribution of both between-subject and within-subject differences (Fig. 4). The absolute difference was averaged for each fingerprint entry to map its spatial distribution. Each voxel can have multiple local connectome fingerprint measurements. For visualization purposes we only calculated the difference of the first resolved fiber (defined by the atlas). The first row of Fig. 4 shows between-subject differences for datasets I, II, III, and IV. The largest between-subject differences are found in core white matter structures such as the corpus callosum and central semiovale. The corpus callosum is known to have commissural fibers connecting the cortical hemisphere, whereas the central semiovale has association pathways connecting frontal, parietal, and occipital regions as well as projection pathways connecting cerebral cortex and brainstem. The large differences found in these two regions suggest that the between-subject differences could be driven by a variety of different brain connections. The second row of Fig. 4 shows within-subject differences for datasets I, II, III, and IV. Dataset I was acquired with the shortest time interval between repeat scans (less than 16 days), whereas dataset II (1∼3 months), dataset III (6 months) and dataset IV (a year) were acquired with longer time intervals. As shown in Fig. 4, the within-subject differences are substantially lower than the between-subject differences, suggesting high uniqueness of the local connectome fingerprint to an individual. Substantial increase of within-subject differences could be observed in the corpus callosum at datasets with a longer time interval, suggesting the possibility of neuroplasticity over time.

**Fig 3.**
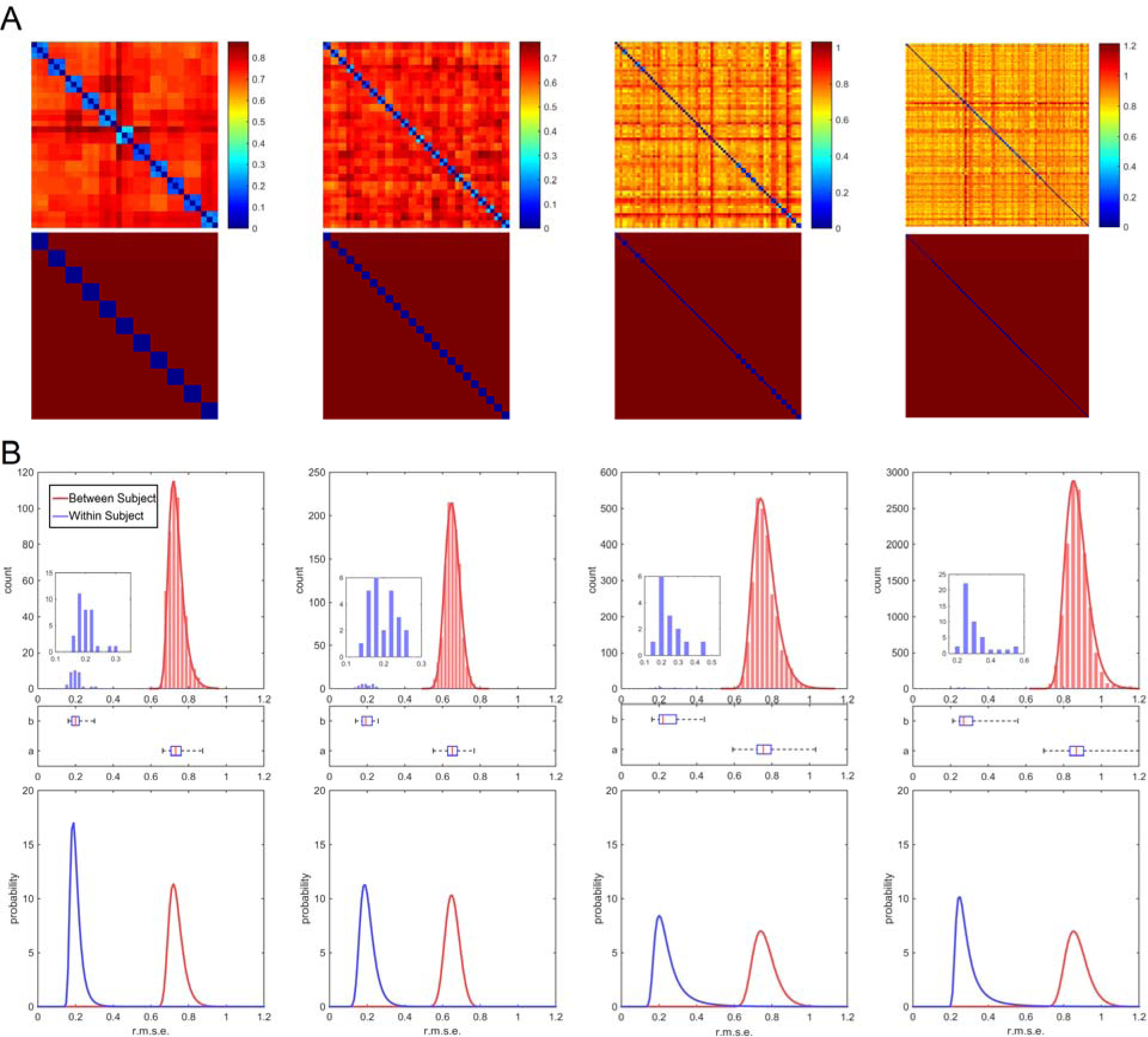
Within versus between-subject differences in the local connectome fingerprints. (A) The first row shows the matrix of pair-wise distances between any two local connectome fingerprints for datasets I, II, III, and IV (column 1, 2, 3, and 4, respectively). The second row shows the location of the within-subject (blue) and between-subject differences (red). (B) The histograms of within-subject (blue) and between-subject (red) differences in the connectome fingerprints calculated from datasets I, II, III, and IV (column 1, 2, 3, and 4, respectively). The first row shows the histograms, and the second row shows the box plot of their quartiles. In these four datasets, within-subject (blue) and between-subject (red) differences have perfect separation. In the last row, the histograms are fitted with generalized extreme value distribution (also shown by solid curves in the second row) to estimate the classification error of the connectome fingerprint. The estimated classification error was 4.25×10^-6^, 9.97×10^-7^, 5.3×10^-3^, and 5.5×10^-3^ for dataset I, II, III, and IV, respectively. The larger error in dataset III and IV could be due to their longer scanning interval (6 months and one year).

**Fig.4.**
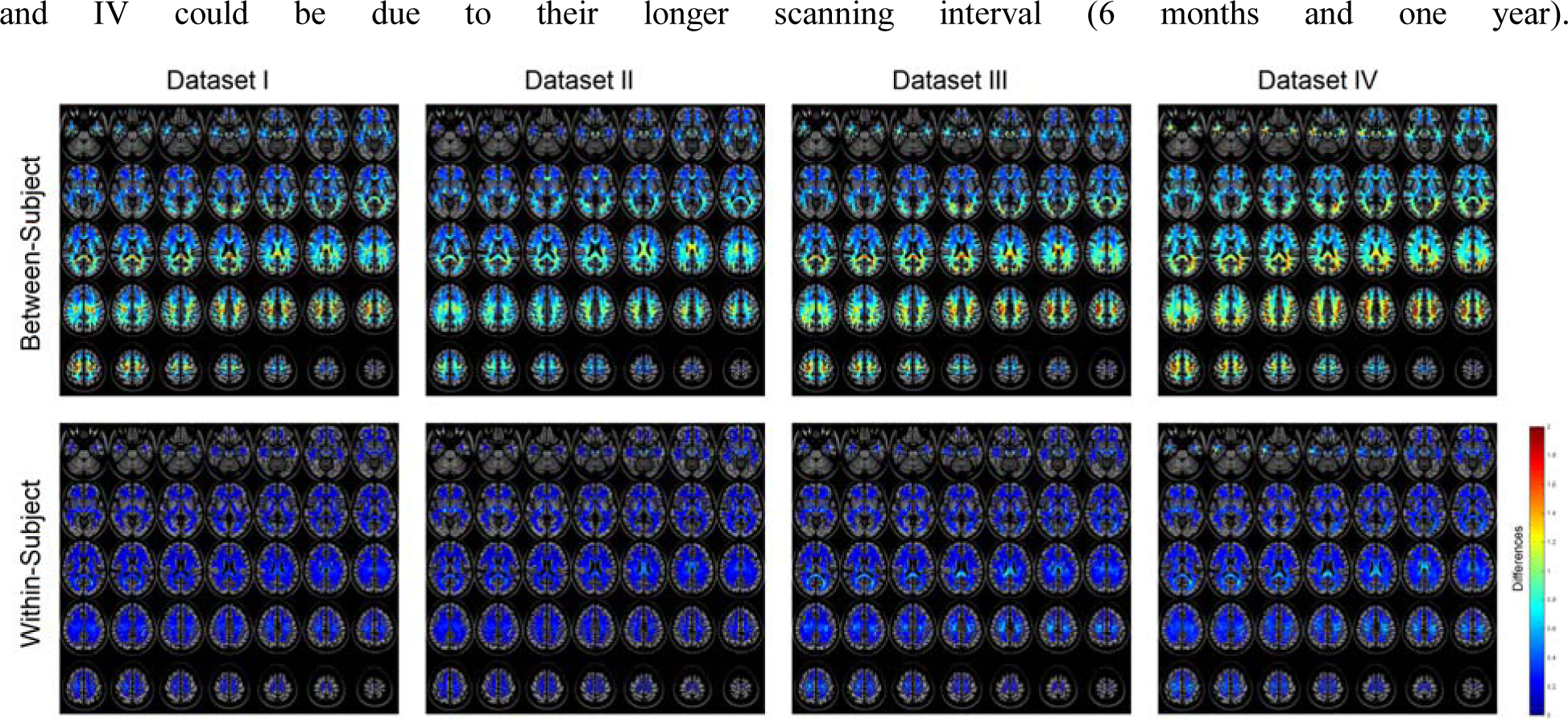
The spatial mapping of between-subject (first row) and within-subject (second row). Dataset I was acquired within 16 days, whereas dataset II (1∼3 months), dataset III (6 months) and dataset IV (a year) were acquired at longer time intervals. High between-subject differences can be observed in white matter tissue, especially the corpus callosum and central semiovale. The within-subject differences are much smaller, and repeat scans with longer time intervals show higher within-subject differences.

This within-subject consistency suggests that the local connectome fingerprint could be used as a unique subject identifier. To assess this, we used a linear discriminant analysis (LDA) classifier [17] to determine whether two fingerprints were from the same individual using only the Euclidian distance between fingerprints as a single classification feature. For each dataset, the classification error was estimated using leave-one-out cross-validation. We did not observe a single misclassification out of the 17,398 cross-validation rounds from four datasets (17,283 different-subject and 115 same-subject pairings). To approximate the classification error, we modeled the distributions of within-subject and between-subject differences by the generalized extreme value distribution [18], a continuous probabilistic function often used to assess the probability of extreme values (smallest or largest) appearing in independent identically distributed random samples (last row of Fig. 3B). The classification error was quantified by the probability of a within-subject difference greater than a between-subject difference. Our analysis showed that the classification error was 4.25×10^-6^ for dataset I, 9.97×10^-7^ for dataset II, 5.3×10^-3^ for dataset III, and 5.5×10^-3^ for dataset IV. The larger error in dataset III and IV could be due to their longer scan interval (6 months and one year). For repeat scans acquired within 3 months, the probability of mistaking two samples of the same subject’s local connectome fingerprint as coming from two different individuals was low enough to consider the local connectome fingerprint a highly reliable measure of individual subject uniqueness.

### Corpus callosum fingerprint

Since gross anatomical patterns such as gyral and sulcal folding can be highly specific to an individual, it is possible that the unique features we observed in the local connectome fingerprint reflected an artifact of the spatial normalization process. To evaluate this, we retested our uniqueness within a restricted white matter mask that only covered the median sagittal sections of the corpus callosum defined by the Johns Hopkins University white matter atlas [19]. This “corpus callosum fingerprint” should be free from all possible contributions of anatomical geometry such as gyral and sulcal folding. We applied the same analysis procedures to the corpus callosum fingerprint to examine whether it can reveal unique patterns specific to individuals within this area. The result showed d-prime values of 5.97, 5.85, 3.78, and 4.08, for dataset I, II, III, and IV, respectively. The leave-one-out cross-validation analysis showed that classification error was 0%, 0.089%, 1.26%, and 0.63%, for dataset I, II, III, and IV, respectively. The classification error modeled by the generalized extreme value distribution was 9.13×10^-4^, 5.6×10^-3^, 6.9×10^-3^, and 7.2×10^-3^, for dataset I, II, III, and IV, respectively. The corpus callosum fingerprint itself already achieved more than 99% accuracy in subject identification. This suggests that the high individuality of the local connectome fingerprint is due to the microscopic characteristics of the white matter architecture.

### Comparison with diffusivity-based fingerprints

Diffusivity-based metrics, such as FA, axial diffusivity (AD), and radial diffusivity (RD), also reveal microscopic structure of white matter systems. To compare these measures against SDF, we used the same analysis and replaced the SDF-based measures with FA, AD, and RD values of the corresponding voxels. Our analysis showed that the d-prime values of the FA-based fingerprint were 4.84, 4.70, 4.56, and 3.60, for dataset I, II, III, and IV, respectively. All values were substantially smaller than the local connectome fingerprint. The leave-one-out cross-validation analysis showed that classification error of the FA-based fingerprint was 0%, 0.18%, 0.22%, and 0.87%. While FA-based fingerprints also have high uniqueness with classification error less than 1%, the performance is not superior to the 0% leave-one-out cross-validation error achieved by the local connectome fingerprint

We also analyzed the performance of AD-based fingerprints, producing d-prime values of 4.20, 4.07, 4.33, and 3.68, for datasets I, II, III, and IV, respectively. The performance was similar to FA-based fingerprints and substantially lower than those of the local connectome fingerprint. The leave-one-out cross-validation analysis showed a classification error of 0.15% in dataset IV. While no misclassification was found in dataset I, II, III, the generalized extreme value distribution showed a classification error of 0.18%, 0.29%, and 0.18%, respectively. The AD-based fingerprint was also inferior to the local connectome fingerprint.

The analysis on RD showed a slightly different result. The d-prime values for RD were 7.87, 9.10, 8.79, and 5.80, which were substantially better than FA-based and AD-based fingerprints. Compared with the local connectome fingerprint, it is noteworthy that the local connectome fingerprint substantially outperformed RD for repeat scans within 3 months (14.84 and 12.80 versus 7.87 and 9.10), but not for repeat scans with a longer time interval (7.21, and 8.12 versus 8.79, and 5.80). The classification error also showed a similar pattern. In dataset I, which had the shortest time interval (less than 16 days), the classification errors were 4.25×10^-6^ for the local connectome fingerprint and 0.28% for the RD-based fingerprint. By contrast, in dataset IV, which had the longest time interval (around a year), the classification errors were 5.5×10^-3^ for the local connectome fingerprint and 3.1×10^-3^ for the RD-based fingerprint. The uniqueness of the local connectome fingerprint dropped substantially over time. These time-dependent differences were further investigated in the *Neuroplasticity revealed by the local connectome fingerprint* section below. To summarize, compared with diffusivity-based fingerprints, the local connectome fingerprints exhibited the greatest reliability for repeat scans acquired within 3 months.

### Comparison with global connectivity-based fingerprints

We further compared the local connectome fingerprint with region-to-region connectivity estimates from diffusion MRI fiber tracking. The same analysis pipeline used for the local connectome fingerprint was used to calculate leave-one-out cross-validation error for the traditional connectivity matrix. The d-prime values for the region-to-region connectivity matrices in dataset I, II, III, and IV were at 3.44, 2.06, 2.41, and 2.25, respectively. The classification error for datasets I, II, III, and IV were 3.6%, 13.65%, 11.81%, and 9.48%, respectively (estimated by leave-one-out cross validation). While the classification accuracy for the traditional connectivity matrices is still quite high and similar to what has previously been observed in resting state functional connectivity estimates [20], it is clear from these results that the greatest reliability at characterizing connectomic uniqueness comes from local connectome measures.

### Neuroplasticity revealed by the local connectome fingerprint

Our analysis of within-subject differences hinted at the possibility of changes in the local connectome over time, and thus we further examined how time impacts the uniqueness of local connectome fingerprints. If the local connectome fingerprint is sensitive to neuroplasticity, a longer interval should result in decreased similarity between repeat scans of the same individual. To test this, we calculated the similarity of within- subject local connectome fingerprints as a percentage of the mean between-subject difference (see Methods). A similarity of 100% indicates that two fingerprints are identical, whereas a similarity of 0% indicates the magnitude of the differences between two fingerprints is the same as those between unrelated subjects.

For this analysis, we calculated the similarity between repeat scans in dataset II (*n*=24), which was acquired with the widest range of time interval between repeat scans (1∼3 months). Fig. 5A shows the scatter plot of the similarity against the time. A nonparametric, rank-based test (the Mann-Kendall test) showed a significant decreasing trend in the similarity over time (*p* = 0.0023). To further quantify the change of similarity in the local connectome fingerprint, we used linear regression to calculate the coefficient (slope) between the time interval and similarity. The results showed that the similarity dropped at a rate of 12.79% per 100 days. It is noteworthy that the identical analysis was applied to FA-based, AD-based, and RD-based fingerprints but none showed a significant trend (*p* = 0.3092, 0.4130, and 0.0702, respectively).

To further illustrate how the local connectome fingerprint revealed neuroplasticity within individuals, we selected one subject in dataset IV that exhibited the greatest difference across time and visualized the spatial mapping of the local connectome fingerprints between repeat scans. This spatial mapping is shown in Fig. 5B, whereas the FA map calculated from the same data are shown in Fig. 5C. The upper row shows the midsagittal view at the corpus callosum, whereas the lower row shows an axial view at the splenium and genu of the corpus callosum. Each voxel may have multiple local connectome fingerprint measurements (e.g. at the crossing fiber region), and for visualization purposes, only the one associated with the first resolved fiber (defined by the atlas) was calculated. All images are scaled by their maximum values to provide a fair comparison. Fig. 5B shows substantial differences in several core white matter bundles between the repeat scan (annotated), whereas Fig. 5C shows no obvious difference. Since artifacts such as signal drift, coil degradation, and motion affect large regions of tissue spanning several centimeters, the fact that the differences in the local connectome fingerprint were observed in specific white matter bundles suggests that the finding is unlikely due to an artifact.

**Fig 5.**
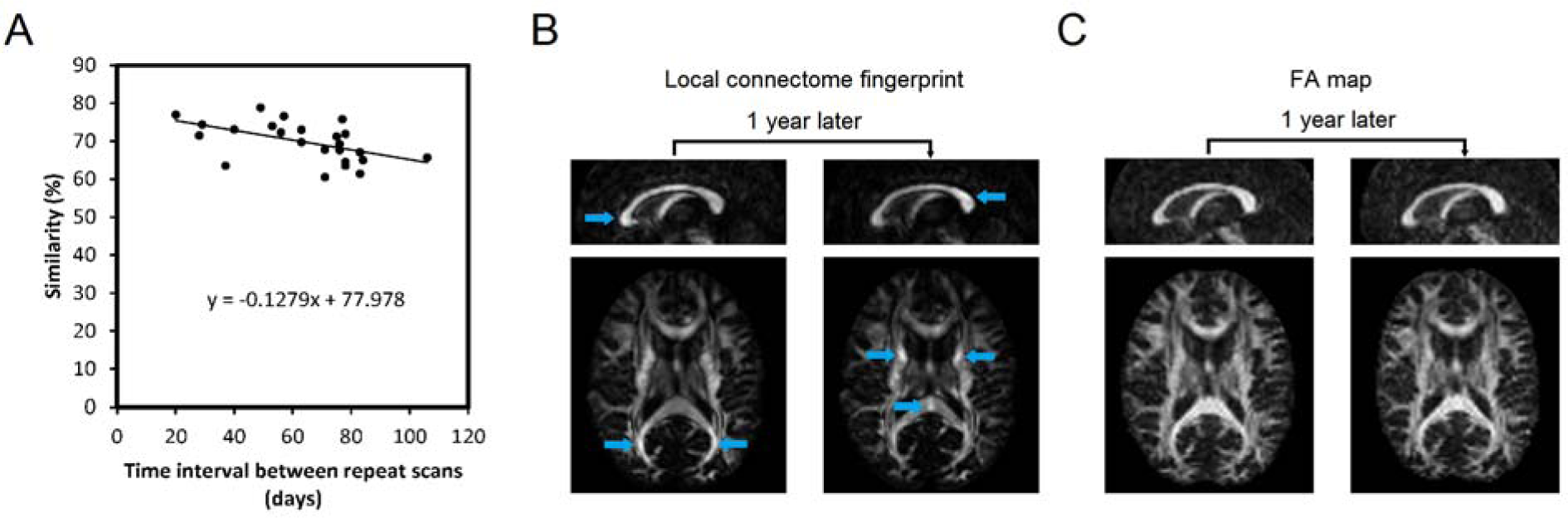
Neuroplasticity revealed by the local connectome fingerprint. (A) A scatter plot showing a decreasing trend in similarity against time in dataset II. (B) The local connectome fingerprint from one subject in dataset IV shows substantial differences between repeat scans. The changes include both increased and decreased connectivity at different locations, leading to a drop in the self-similarity. The fact that these changes are located at specific white matter bundles suggests that they are unlikely due to an image artifact. (C) The FA map calculated from the same data shows no obvious difference between repeat scans. There is no visible trait of signal drift, motion artifact or coil degradation, confirming the quality of the image acquisition.

### Similarity among genetically-related individuals

The local connectome fingerprint opens the possibility for comparing not only differences but also the similarities between individuals. To further illustrate how the local connectome fingerprint can be used to quantify white matter architecture as a phenotypic marker, we used a publicly available dMRI dataset of 486 subjects from Human Connectome Project (2014, Q3 release), including 49 pairs of monozygotic (MZ) twins, 43 pairs of dizygotic twins (DZ) twins, and 96 pairs of non-twin siblings. While the local connectome fingerprints of MZ twins show generally similar patterns at the coarse level (Fig. 6), there are also substantial individual differences between the twins that can be observed along the fingerprints. Consistent with these qualitative comparisons, we found that MZ twins have smaller differences between fingerprints, followed by DZ twins, siblings, and unrelated subjects (Fig. 7A). It is noteworthy that all difference distributions have large overlapping regions (Fig. 7B), indicating that the difference between twins or siblings may often fall within the distribution of differences from genetically-unrelated subjects. We further compared the similarity between twins and siblings. On average, MZ twins have a similarity index of 12.51±1.09%, whereas similarity for DZ twins and siblings is 5.14±1.34% and 4.47±0.59%, respectively (Fig. 7C). The difference in similarity index was significant across MZ twins, DZ twins, non-twin siblings, and other genetically-unrelated subjects (Kruskal-Wallis test, χ^2^[3,22895] = 165.43, *p* < 0.001). Post-hoc comparisons using Scheffé's S procedure showed (1) significantly higher similarity in MZ twins compared with all other groups (all *p* < 0.001), (2) significantly higher similarity in DZ twins compared with unrelated subjects (*p* = 0.001), and (3) significantly higher similarity in non-twin siblings compared with unrelated subjects (p = 0.0146). There was no significant difference between DZ twins and non-twin siblings ( *p*= 0.9989). This result is consistent with MZ twin sharing a higher genetic similarity, whereas DZ twins exhibit a similar genetic similarity on par with non-twin siblings.

**Fig 6.**
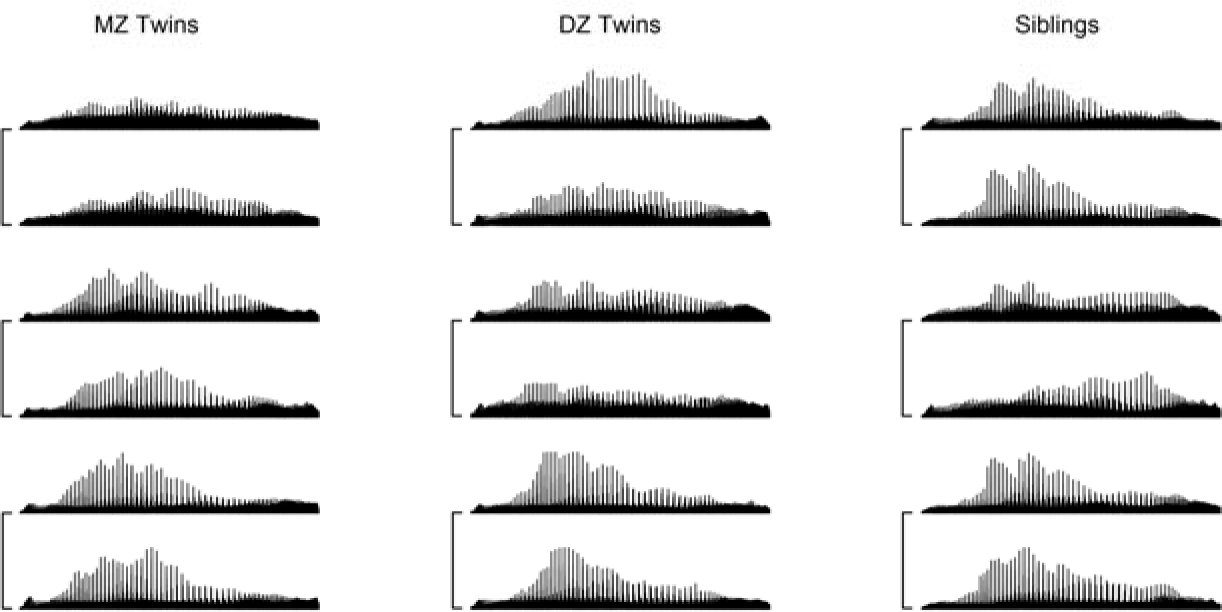
The local connectome fingerprints of monozygotic (MZ) twins, dizygotic (DZ) twins, and non-twin siblings. Three pairs of connectome fingerprints are shown for each group, Pairs are annotated by a connecting line. The connectome fingerprints between MZ twins show the grossly similar patterns though some between-subject difference can still be observed. DZ twins and siblings also have a similar pattern, but the between-subject difference becomes more prominent.

**Fig 7.**
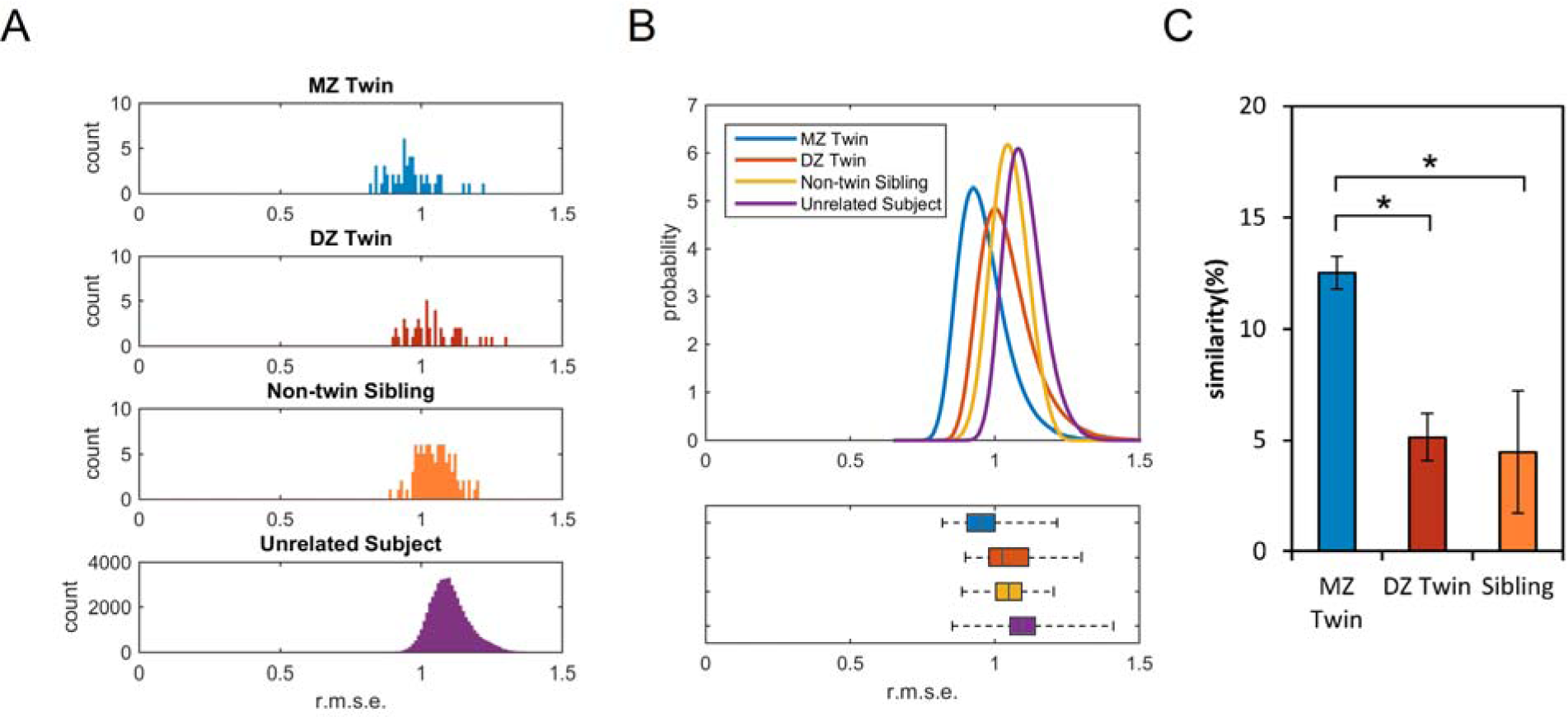
The differences and similarities between twins and siblings. (A) The histograms show the distribution of the root-mean-square-error (r.m.s.e) between MZ twins, DZ twins, non-twin siblings, and genetically unrelated subjects calculated from their local connectome fingerprints. On average, MZ twins have the lowest difference between each twin pair, followed by DZ twins and siblings. (B) The upper figure shows the differences fitted with generalized extreme value distribution. The lower figure shows the box plot of the distribution to facilitate comparison. The four distributions are mostly overlapping, indicating that twins and siblings still have high individuality similar to genetically-unrelated subjects. (C) The similarity between MZ twins is significantly higher than that between DZ twins or non-twin siblings, whereas the similarity between DZ twins is not statistically different from the similarity between non-twin siblings.

## Discussion

Local white matter architecture is so unique and highly conserved within an individual that it can be considered a unique neural phenotype. Here we show that this phenotype can be quantified by measuring the density of microscopic water diffusion along major white matter fascicles and producing a high dimensional vector that can be used to compute the distance between two structural connectomes, i.e., a local connectome fingerprint. The distance between two local connectome fingerprints reflects a low dimensional representation of both similarities and differences in whole-brain white matter pathways. Our analysis showed how the local connectome fingerprint exhibited unprecedentedly high between-subject distance, while generally low within-subject distances, allowing for it to be used as a reliable measure of the specific connective architecture of individual brains. This property paves the way for using the local connectome as a phenotypic marker of the structural connectome.

The concept of local connectome is both conceptually and methodologically different from conventional connectomic measures. While most studies have emphasized on region-to-region connectivity [3] and ignored the rich information in the local white matter architecture, the local connectome reveals the connectivity at the voxel level and characterizes local white matter architecture to provide high dimensional data that may complement the region-to-region connectivity [13]. This local connectome mindset considers the fact that the difference between brain structures may be localized and thus may not be readily identified in the global connectomic pattern. We have previously shown that the local connectome can be used to localize the change of white matter structure due to physiological difference such as body mass index [13].

While any high dimensional representation of the human brain in a standard space has the potential to be used as fingerprint, we showed that the uniqueness of fingerprints generated from the local connectome was substantially higher than what was observed in diffusivity-based fingerprints as well as fingerprints derived from region-to-region connectivity reported by either dMRI or fMRI, as typically done in human connectomic studies [2, 20, 21]. For example, the region-to-region structural connectivity achieved a classification accuracy around 90∼97%. This is very close to the accuracy of its functional counterpart [20], that was recently reported to have an accuracy of 92-94% in whole brain identification and 98-99% in frontoparietal network. Although both region-to-region connectivity approaches have accuracy greater than 90%, the performance remains substantially lower than the perfect classification in 17,398 leave-one-out rounds and an estimated error of 10^-6^ achieved by local connectome fingerprint.

At first glance, it may seem possible that the high degree of uniqueness exhibited by the local connectome fingerprint could be due to variability in the spatial normalization process between individuals driven by the unique gyral or sulcal folding patterns in gray matter. While we still cannot rule out the effect of misalignment, our comparison with the FA-based fingerprints showed that the spatial normalization process does not fully contribute to the uniqueness observed in the local connectome fingerprint. Both FA-based fingerprints and the local connectome fingerprints used an identical spatial normalization mapping process, but the FA-based fingerprints had a much higher error rate in leave-one-out cross-validation (e.g. 0.87% for dataset IV) than the zero cross-validation error achieved by the local connectome fingerprint. Obviously, a substantial portion of the uniqueness was due to the microstructural white matter characteristics quantified in the SDF. Moreover, we observed favorable characterization of white matter uniqueness even when our analysis was restricted to a small portion of white matter with minimal influence of sulcal and gyral folding (i.e., the mid corpus callosum). These two findings support our claim that the local connectome fingerprint can reveal the unique characteristics of the white matter architecture. Finally, the between-subject differences are mostly located within the deep white matter at the central semiovale and the corpus callosum. This spatial specificity suggests that the uniqueness of the local connectome fingerprint is mostly driven by mesoscopic or microscopic architectural properties, not due to an artifact of unique folding geometry or the spatial normalization process.

It is important to point out that the local connectome fingerprint is based on a physical measurement that is different from diffusivity-based metrics such as FA, AD, and RD. To further compare their physical meanings, diffusivity quantifies *how fast* water diffuses in tissue [22] and is sensitive to the structural integrity of the underlying fiber bundles [14], such as axonal loss and demyelination [23-26]. This may explain why the FA map appears similar across the normal population in which the axons have normal structure. By contrast, SDF quantifies *how much* water diffuses along the fiber pathways [15, 27] and is sensitive to density characteristics of white matter such as the compactness of the fiber bundles [15, 28, 29]. As illustrated in our qualitative analysis (Fig. 1C), while the density characteristics vary substantially among normal populations, the FA measurements do not show obvious differences between subjects. This highlights how the local connectome fingerprint achieved a higher uniqueness profile than diffusivity-based metrics when they were used to characterize microstructural white matter patterns that reflect individuality. The results led us to hypothesize that the local connectome fingerprints may be more sensitive to axonal density or different levels of myelination that are unique to individuals. Future histology studies are needed to confirm this hypothesis.

The high degree of uniqueness in the local connectome within an individual can be used to reflect a quantifiable phenotype of neural organization. As illustrated in our analysis of twins, the similarity in monozygotic twins was around twice as much of the dizygotic twins, whereas our post-hoc analysis did not find significant similarity difference between dizygotic twins and non-twins siblings. These results are highly suggestive that genetics contribute a substantial portion to the overall construction of the local connectome, which is consistent with previous studies showing high heritability in cortical connections [30, 31] and white matter integrity [32-35]. Nevertheless, our results also showed that the monozygotic twins shared only 12.51% similarity in local white matter architecture. This indicates that a high heritability may not necessarily imply that most of the differences or similarity observed in phenotypes are due to genetic factors [36]. A considerable portion of the individuality in local connectome is likely driven by environmental factors such as life experience and learning. Thus monozygotic twins still exhibited high individuality in their connectome. In fact, our findings showed that the local connectome fingerprint is highly plastic over time, presented by a significant decreasing trend in the self-similarity caused by either an increase or decrease in the local connectome fingerprint measurements. This decreasing trend in the self-similarity raises many questions about which factors (genomic, social, environmental, or pathological) sculpt the local white matter systems. Of course, white matter integrity also varies with normative development [37-39], a portion of which may be determined genetically. This warrants more longitudinal and genetic analysis to identify specific contributions of genetic and environmental factors on the uniqueness of connectomic structure, with an aim to understand how those factors interact with abnormal brain circuits in neurological and psychiatric disorders.

It is important to point out that the highest similarity between repeat scans was around 70∼80% in our study. This indicates that 20-30% of variability in the local connectome may arise from artifacts that decrease signal-to-noise ratio, such as cardiovascular and respiratory artifacts or computation error. This number reflects the limit of the local connectome fingerprint in detecting an anomaly in the individuals as well as finding differences in a group study. For example, we could not accurately identify whether two scans were from a twin pair because the similarity between twins was only around 12.51%. However, if a disease causes a white matter change with more than 30% difference in similarity, the local connectome fingerprint may be able to detect it. In a group study, increasing the number of subjects can average out the effect of noise and error on the similarity, allowing us to find a group difference that is substantially small. The similarity index from repeat scans allows us to gauge the strength and limitation of the local connectome fingerprint and prospectively, to develop a strategy to improve its performance.

## Methods

### Five independently collected dMRI datasets

The first dataset included a total of 11 subjects (9 males and 2 females, age 20∼42). Each subject had three diffusion MRI scans within 16 days on a Siemens Trio 3T system at the University of California, Santa Barbara. All methods were approved by the local institutional review board at the University of California, Santa Barbara. The diffusion MRI was acquired using a twice-refocused spin-echo EPI sequence. A 257-direction full-sphere grid sampling scheme was used. The maximum b-value was 5000 s/mm^2^. TR = 9916 ms, TE = 157 ms, voxel size = 2.4×2.4×2.4 mm, FoV = 231×231 mm.

The second set of data included a total of 24 subjects (8 males and 16 females, age 22 ∼ 74). All participants were scanned on a Siemens Tim Trio 3T system at National Taiwan University, and all subjects had their second scan at 1∼3 months. All methods were approved by the local institutional review board at National Taiwan University. The diffusion MRI was also acquired using a twice-refocused spin-echo EPI sequence. The diffusion scheme is a 101-direction half-sphere grid sampling scheme with b-max = 4000 s/mm^2^ (b-table available at http://dsistudio.labsolver.org). TR = 9600 ms, TE = 130 ms, voxel size = 2.5×2.5×2.5 mm.

The third set of data included a total of 60 subjects (30 males and 30 females, age 18 ∼ 46). All participants were scanned on a Siemens Verio 3T system at Carnegie Mellon University, and 14 of the 60 subjects had their second scan at 6 months. All methods were approved by the local institutional review board at Carnegie Mellon University. The diffusion MRI was also acquired using a twice-refocused spin-echo EPI sequence. A 257-direction full-sphere grid sampling scheme was used. The maximum b-value was 5000 s/mm^2^. TR = 9916 ms, TE = 157 ms, voxel size = 2.4×2.4×2.4 mm, FoV = 231×231 mm.

The fourth set of diffusion data included a total of 118 subjects (91 males and 27 females, age 22 ∼ 55) that were also scanned on a Siemens Verio 3T system at the Carnegie Mellon University. All methods were approved by the local institutional review board at the University of Pittsburgh and Carnegie Mellon University. 44 of them had another scan after one year. The diffusion images were acquired on a Siemens Verio scanner using a 2D EPI diffusion sequence. TE=96 ms, and TR=11100 ms. A total of 50 diffusion sampling directions were acquired. The b-value was 2000 s/mm^2^. The in-plane resolution was 2.4 mm. The slice thickness was 2.4 mm.

The fifth dataset was from the Human Connectome Projects (Q3, 2014) acquired by Washington University in Saint Louis and University of Minnesota. The diffusion MRI data were acquired on a Siemens 3T Skyra scanner using a 2D spin-echo single-shot multiband EPI sequence with a multi-band factor of 3 and monopolar gradient pulse. A total of 486 subjects (195 males and 291 females, age 22 ∼ 36) received diffusion scans. The spatial resolution was 1.25 mm isotropic. TR=5500 ms, TE=89.50 ms. The b-values were 1000, 2000, and 3000 s/mm^2^. The total number of diffusion sampling directions was 90, 90, and 90 for each of the shells in addition to 6 b0 images. The total scanning time was approximately 55 minutes. The scan data included 49 pairs of monozygotic twin, 43 pairs of dizygotic twins, and 96 pairs of siblings. We used the preprocessed data provided by the consortium in our analysis. Carnegie Mellon University Institutional Review Board (IRB) reviewed the research protocol for the data analysis in accordance with 45 CFR 46 and CMU’s Federal-wide Assurance. The research protocol has been given approval as Exempt by the IRB on March 12, 2014, in accordance with 45 CFR 46.101(b)(4) (IRB Protocol Number: HS14-139).

### Local connectome fingerprinting

All five datasets were processed using an identical processing pipeline implemented in DSI Studio (http://dsi-studio.labslover.org), an open-source diffusion MRI analysis tool for connectome analysis. The source code is publicly available on the same website. As shown in Fig. 2A, the diffusion MRI data of each subject were reconstructed in a common stereotaxic space using q-space diffeomorphic reconstruction (QSDR)[40], a white matter based nonlinear registration approach that directly reconstructed diffusion information in a standard space:

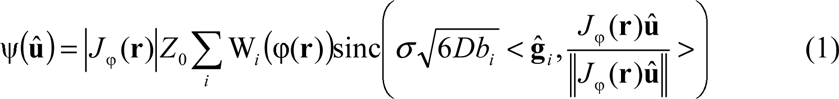

ψ(**û**) is a spin distribution function (SDF)[15] in the standard space, defined as the density of diffusing spins that have displacement oriented at direction **û**. φ is a function that maps a coordinate **r** from the standard space to the subject’s space, whereas *J*_φ_ is the Jacobian matrix of φ, and |*J*_φ_| is the Jacobian determinant. W_*i*_ is the diffusion signals acquired by a b-value of *b*_*i*_ with diffusion sensitization gradient oriented at ĝ_*i*_. σ is the diffusion sampling ratio controlling the displacement range of the diffusing spins sampled by the SDFs. Lower values allow for quantifying more from restricted diffusion. *D* is the diffusivity of free water diffusion, and *Z*_0_ is the constant estimated by the diffusion signals of free water diffusion in the brain ventricle [40]. A σ of 1.25 was used to calculate the SDFs, and 1 mm resolution was assigned to the output resolution of the QSDR reconstruction.

A common axonal directions atlas, derived from the HCP dataset (this HCP-488 atlas is freely available at http://dsi-studio.labsolver.org), was used as a common SDF sampling framework to provide a consistent set of sampling directions **û** to sample the magnitude of SDFs along axonal directions in the cerebral white matter. Gray matter was excluded using the ICBM-152 white matter mask (MacConnel Brain Imaging Centre, McGill University, Canada). The cerebellum was also excluded due to different slice coverage in cerebellum across subjects. Since each voxel in the cerebral white matter may have more than one axonal direction, multiple measurements can be extracted from the SDF of the same voxel. The density measurements were sampled by the left-posterior-superior voxel order and compiled into a sequence of scalar values (Fig. 2B). Since the density measurement has arbitrary units, the local connectome fingerprint was scaled to make the variance equal to 1.

### Estimation of classification error

For each dMRI dataset, the root-mean-squared error between any two connectome fingerprints was calculated to obtain a matrix of paired-wise difference. The calculated difference was used as the feature to classify whether two connectome fingerprints are from the same or different person. The default linear discriminant analysis (LDA) classifier provided in MATLAB (MathWorks, Natick, MA) was used, and for each dataset, the classification error was estimated using leave-one-out cross-validation. We also used a modeling method to calculate the classification error if the leave-one-out cross-validation did not yield any classification error. The histograms of the within-subject and between-subject differences were fitted by the generalized extreme value distribution using the maximum likelihood estimator (gevfit) provided in MATLAB. To consider the non-negativity of the distribution, the estimated *k* parameter of the generalized extreme value distribution was set to be greater than 0. The classification error was estimated by the probability of a within-subject difference greater than a between-subject difference estimated using the generalized extreme value distribution.

### Comparison with traditional connectivity matrix

To compare local connectome fingerprint with region-to-region connectivity matrix, deterministic fiber tracking[28] was applied using a 100,000 uniform white matter seeding points, a maximum turning angle of 60 degrees, and a default anisotropy threshold determined using Otsu’s threshold [41]. The cortical regions were defined through a nonlinear registration between the subject anisotropy map and the HCP-488 anisotropy map in DSI Studio and parcellated using the Automated Anatomical Labeling (AAL) atlas. The matrix entries were quantified by the number of tracks ending in each of the region pairs. The root-mean-squared error can also be calculated from any two connectivity matrices. The classification error was also estimated and compared with local connectome fingerprint.

### Similarity index

The similarity index between two local connectome fingerprints was calculated by 100%×(1-d_1_/d_0_), where d_1_ was the difference between two fingerprints, and d_0_ was the expected value of the differences between unrelated subjects scanned by the same imaging protocol. The similarity between MZ twins, DZ twins, non-twin siblings, and repeated scans was calculated and compared. The Kruskal-Wallis test was applied to four groups (MZ and DZ twins, siblings, and unrelated subjects) with independent samples. To further study the similarity between repeat scans, the similarity indices were tested against their scanning time intervals by the Mann-Kendall test to study the effect of time interval on the local connectome fingerprints.

## Acknowledgments

The research was sponsored by the Army Research Laboratory and accomplished under Cooperative Agreement Number W911NF-10-2-0022. The views and conclusions contained in this document are those of the authors and should not be interpreted as representing the official policies, either expressed or implied, of the Army Research Laboratory or the U.S. Government. This research was supported by and NSF BIGDATA (1247658). Part of the data used in this study were from the Human Connectome Project, WU-Minn Consortium (1U54MH091657). This research was supported in part by the Ruentex Group and the Ministry of Economic Affairs, Taiwan (101-EC-17-A-19-S1-175). This research was supported in part by National Institutes of Health (R01 DK095172).

